# Sequence, Structure, and Epitope Analysis of the Polymorphic Membrane Protein Family in *Chlamydia trachomatis*

**DOI:** 10.1101/2022.07.13.499933

**Authors:** Patrick W. Cervantes, Brent Segelke, Edmond Y. Lau, Luis de la Maza, Matthew Coleman, Patrik D’haeseleer

## Abstract

The polymorphic membrane proteins are a family of autotransporters that play an important role in infection, adhesion and immunity in *Chlamydia trachomatis*. Here we show that the characteristic GGA(I,L,V) and FxxN tetrapeptide repeats fit into a larger repeat sequence, and that these repeats correspond to the coils of a large beta-helical domain in high quality structure predictions. While the tetranucleotide motifs themselves are predicted to play a structural role in folding and close stacking of the beta-helical backbone of the passenger domain, we found many of the interesting features of Pmps are localized to the side loops jutting out from the beta helix - including protease cleavage, host cell adhesion, and B-cell epitopes; while T-cell epitopes are predominantly found in the beta-helix itself.

## Introduction

*Chlamydia trachomatis* is the most common sexually transmitted bacterial infection in humans with an estimated prevalence of 1-2% in the United States and 4.2% globally (1). However, the frequency of *C. trachomatis* infection in humans is likely much higher since nearly 50% of male and 80% of female infections are asymptomatic (1). Untreated *C. trachomatis* infection poses a significant health risk to humans, resulting in blindness, pelvic inflammatory disease, ectopic pregnancy, and infertility (1). There are 15 main *C. trachomatis* genotypes or serovars that are classified into 3 groups based on disease outcomes. These include ocular trachoma (serovars A, B, Ba, C), urogenital infection (serovars D, E, F, G, H, I, J, K), and lymphogranuloma venereum (LVG) (serovars L1, L2, L3). Serovars are classified based on major outer membrane protein (Momp) sequence diversity and immunotyping (2,3). Of the different *C. trachomatis* serovars, E and D are the most prevalent in humans (cite). According to the Center for Disease Control, human clinical manifestations of *C. trachomatis* translated to $144 million in direct medical costs in the US alone in 2018. Treatment options rely on antibiotics for known exposures or symptomatic infection, but the high prevalence of asymptomatic infection necessitates the development of protective vaccines, which is expertly reviewed by de la Maza et al. (2021) (1). In short, vaccines that target Chlamydia surface proteins show promise to elicit protection, decrease transmission, and prevent adverse health outcomes.

The polymorphic membrane protein (Pmp) family is one group of surface-exposed proteins in *Chlamydia* that are viable vaccine candidates (4–8). Several laboratories have shown Pmps are immunogenic in naturally infected humans, as well as in non-human primates and mice infected with *C. trachomatis* or *C. muridarum*, respectively (4,9–12). These studies showed, subunit vaccinations with Pmp N-terminal domains were more immunogenic and suggest cross-serovar and cross-species protection against *Chlamydia*. To further support the potential use and development of a Pmp vaccine, the N-terminal domain was shown to contain novel T-cell epitopes that neutralize a vaginal Chlamydia challenge (13)(14)(15).

Pmps are part of the Type-Va secretion system in gram-negative bacteria that contain an autotransporter domain (16,17). Autotransporters are characterized by 3 functional domains: (i) a secretory sequence that facilitates transport across the plasma membrane, (ii) an outer membrane β-barrel transmembrane region, and (iii) an extracellular passenger domain that carries out a number of biological roles, from enzymatic processes to pathogenesis (18). The Pmp family is further defined by FxxN and GGA (I,L,V) tetrapeptide repeats that are concentrated in the extracellular passenger domain. These motifs occur 13.6 and 6.5 times on average in *C. trachomatis*, respectively, and are thought to play a role in cellular adhesion and infectivity (4,16,19–23). However, it remains unclear whether the tetrapeptide motifs directly mediate cellular adhesion or provide passenger domain structural support.

The number of *pmp* genes vary among *Chlamydia* species. In *C. trachomatis* there are 9 *pmp* genes, all of which are known to be transcribed and translocated to the outer membrane (24–26). Within a *C. trachomatis* serovar, Pmp amino acid sequences collectively have a 50% maximum identity (16). Whereas between serovars, individual Pmps are highly conserved between serovars with Pmps A and D being the most conserved (~98%) and Pmp F the least conserved (~85%) (27,28). Interestingly, phylogenetic analysis and sequence diversity of Pmp H and F, unlike Momp, groups *C. trachomatis* serovars into 3 clades that correspond to ocular, genital, and LGV disease types (27,29), This suggests Pmp’s could play a role in tissue specific infection.

The passenger domain was predicted to form a right-handed β-helix structure (what method?) (30,31). Although there is no crystal structure to validate the computational model of the passenger domain β-helix or the overall Pmp native structure. *C. trachomatis* Pmp passenger domains have been shown to form hetero-and homo-meric oligomers *in vitro* (22,32,33). In addition, proteolytic cleavage sites are predicted for all *C. trachomatis* Pmps, but experimentally validated for only two Pmp orthologs, Pmp D of *C. trachomatis* and Pmp 21 of *C. pneumonia* (21,25,31,33–36). These studies give rise to the idea that Pmps could form diverse oligomeric structures that could contribute antigenic diversity and immune evasion.

In this study we utilize protein structure prediction algorithms to visualize, for the first time, the Pmp family from *C. trachomatis* Serovar E. In addition, our Pmp amino acid sequence analysis discovered the tetrapeptide motifs,GGA(I,L,V) and FxxN, fit into a larger and predictable spacing pattern. Our sequence and structural analysis more accurately defines the Chlamydia’s Pmp family which could be used to inform rational vaccine design and functional studies.

## Methods

### Identifying Pmp repeats

The Multiple Alignment using Fast Fourier Transform (MAFFT) tool was used to align 34 Pmp amino acid sequences. The Pmp amino acid sequences used included all 9 Pmps from *C. trachomatis* Serovar E (Bour), all 9 *C. muridarum* Pmps, and 16 *C. pneumoniae* Pmps.

### Protein structure prediction

After trimming off signal peptides, protein sequences for all nine *C. trachomatis* serovar E Pmps, PmpD of *C. trachomatis* serovar L2, and Pmp21 of C. pneumoniae were submitted for protein structure prediction using both the TrRosetta (37) and RoseTTAFold (38) algorithms through the Robetta protein structure prediction service hosted by the Baker lab (https://robetta.bakerlab.org/). The five models generated by Robetta for each sequence were inspected visually and a representative model chosen. For proteins longer than 1000aa (the length limit on the Robetta server at the time these jobs were submitted), the first and last 1000aa were submitted for structure prediction separately. The resulting models were then aligned using Matchmaker in UCSF Chimera (39), and representative models with good structural homology spliced together into a single structure. AlphaFold structure predictions for the Serovar D homologs of these proteins have recently been released as well, available through Uniprot or directly through the AlphaFold Protein Structure Database at https://alphafold.ebi.ac.uk/faq (40,41)

### Prediction of membrane binding domains

We used the DREAMM public web server at https://dreamm.ni4os.eu/ to predict the membrane-penetrating amino acids in Pmp proteins. DREAMM is a recently developed ensemble machine learning algorithm for predicting protein–membrane interfaces of peripheral membrane proteins (42).

### Molecular dynamics

All molecular dynamics simulations utilized the MARTINI coarse-grain force field (version 2, the proteins used the version 2.2 parameters) (43,44). The system was prepared using the CHARMM-GUI (45). Each protein was embedded into a POPC bilayer, solvated with the MARTINI polarizable water model (46), and the appropriate number of Na + /Cl - was added to neutralize the system and obtain a concentration of 150 mM. The program GROMACS (version 2021.1) was used for the MD simulations (47). The system was energy minimized and 1 microsecond of production dynamics was performed. Simulations were performed in the NPT ensemble with weak temperature and pressure coupling using a velocity-rescaling thermostat at 303 K (coupling constant 1.0 ps) (48) and the Parrinello-Rahman barostat (coupling constant 12 ps) (49). A timestep of 20 fs was used. Electrostatics were calculated with a reaction field (dielectric constant of 2.5) and cutoff at 1.1 nm. The van der Waals interactions were calculated with a cutoff at 1,1 nm.

### B-cell epitope prediction

The full length *C. trachomatis* Serovar E (Bour) Pmp D amino acid sequence was used for B-cell epitope prediction through the Immune epitope database (IEDB) web server (50). The B-cell epitope predictions were generated with BepiPred-2.0 (51) and Discotope (52), two tools that rely on sequence and structure based prediction methods, respectively.

### T-cell epitope prediction

For prediction of MHC Class I and II T-cell epitopes, we used the prediction tools available through the Immune epitope database (IEDB) web server (50), using their recommended settings, which currently default to NetMHCPan 4.1 EL for MHC-I, and Consensus 2.22 for MHC-II.

### Pmp amino acid sequence conservation

Multiple Alignment using Fast Fourier Transform (MAFFT, v7.487) (53) was used to align *C. trachomatis* Pmp amino acid sequences from 15 serovars (A-K, Ba, and L1-L3). Alignments were uploaded to the Multalign View tool in UCSF Chimera and the percent amino acid conservations were mapped onto the respective *C. trachomatis* Serovar E Pmp structural model generated in RoseTTAFold.

## Results

### Pmp passenger domains contain a larger repeat sequence that incorporates the GGA(I,L,V) and FxxN repeat motifs

To test the statement by Grimwood and Stephens that “the number of amino acids between the [GGA(I,L,V) and FxxN] repeat motifs does not appear to follow any characteristic spacing pattern” (16), we examined the spacing between motifs in all nine *C. trachomatis* serovar D and *C. muridarum* Pmps, and all 16 *C.pneumoniae* Pmps using an HMM model. We found that GGA(I,L,V) motifs are almost always followed by a FxxN motif (Figure 2A), and 75% of these pairs (169/226) are spaced only 14-18 amino acids apart (Figure 2B). The spacing following FxxN motifs is much more variable. In about one third of the cases (122 instances), an FxxN motif is followed by a GGA(I,L,V) motif with a gap of only 4-5aa, but much longer gaps or a second FxxN motif are also common. Out of the 34 Pmp proteins examined, we found 188 instances of the core pattern G-G-A-[ILV]-x(14,18)-F-x-x-N, and 301 instances if we allowed one amino acid difference to the core pattern, or allowed the gap to vary from 12-23 amino acids. Figure 2C shows the sequence logo for the HMM model derived from these sequences, showing several conserved amino acid positions between the classical tetrapeptide motifs, including an asparagine at position 15 conserved in 55% of the sequences, and alternating hydrophobic amino acids at positions 4, 6, 10,12, 19, 21, which are likely facing toward the inside of the β-helix.

**Figure 1:**
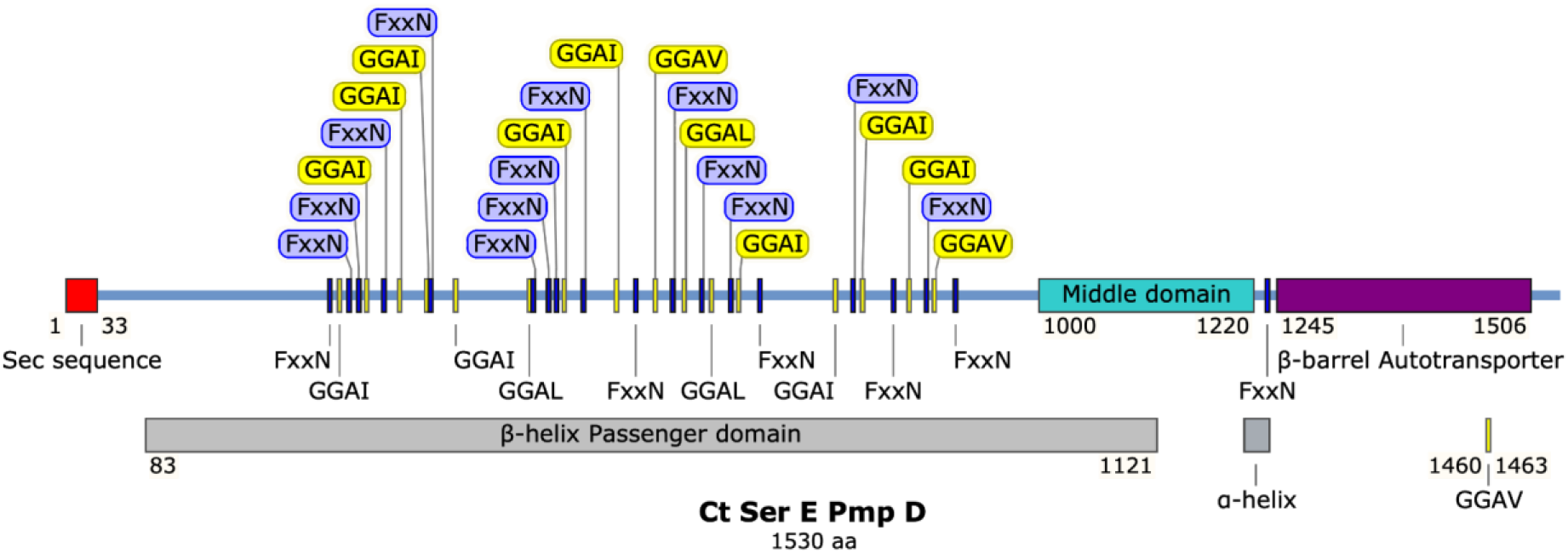
*C. trachomatis* Serovar E (Bour) PmpD protein map. The total PmpD protein length is 1530 amino acids. The protein map shows functional domains which include a secretory sequence, the extracellular β-helix passenger domain, the middle domain, and the transmembrane β-barrel autotransporter. The FxxN and GGA (ILV) tetrapeptide motifs are shown in purple and yellow, respectively. The motif repeats are concentrated in the extracellular passenger domain but can also be found, less frequently, in the C-terminal domain.

**Figure 2:**
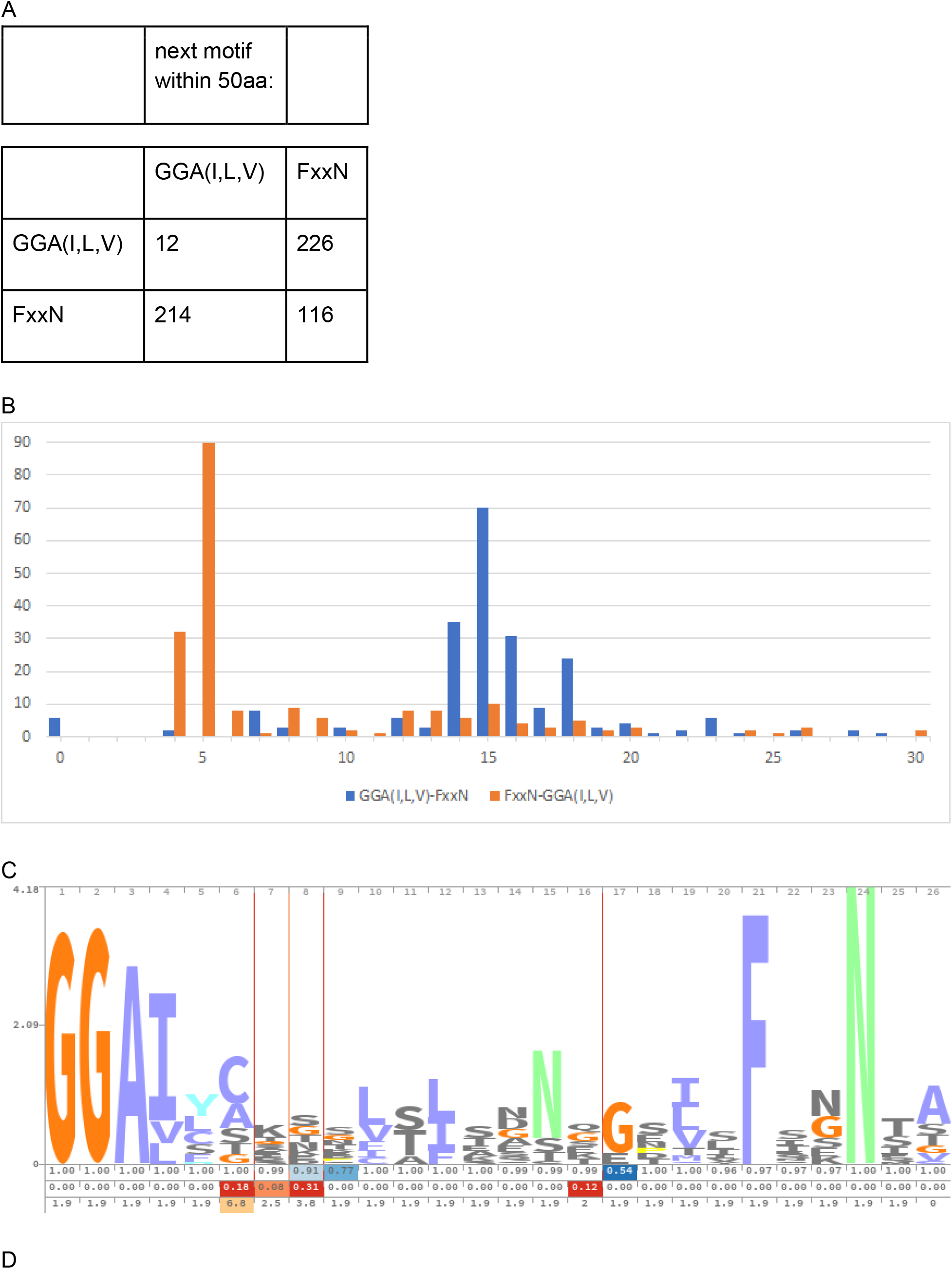
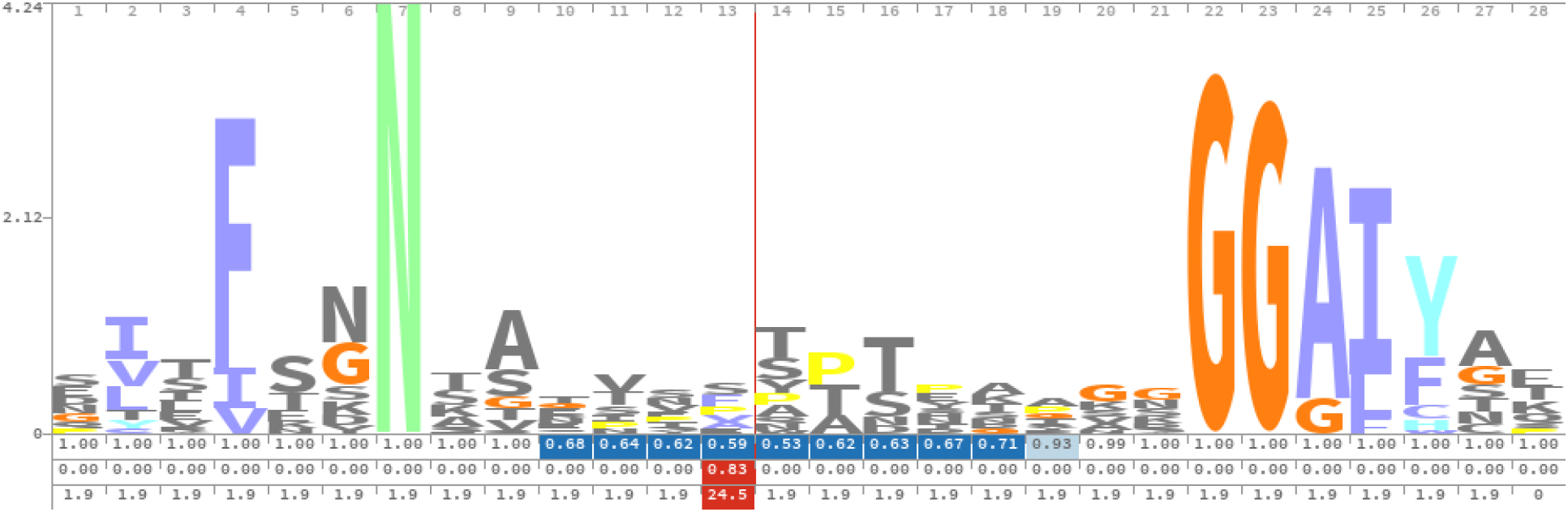
HMM model shows Pmp tetrapeptide repeats, GGA(I,L,V) and FxxN, fit in a larger repeat. (A) The GGA(I,L,V) and FxxN motifs tend to alternate, with GGA(I,L,V) almost always followed by FxxN. (B) GGA - FxxN repeats show a regular spacing. (C) Sequence logo for longer repeat we identified. (D) Sequence logo for PF02415.

Interestingly, Pfam contains a Chlamydia polymorphic membrane protein (Chlamydia_Pmp) repeat domain (PF02415) which incorporates both tetrapeptide motifs, but in the opposite order: FxxN, followed by a long and variable gap with little sequence specificity, followed by GGA(I,L,V) (Figure 2D), missing many of the conserved positions we identified. Although this is supposedly a Chlamydia specific Pfam domain, less than a third of the proteins containing this domain −443 out of 1448 Uniprot proteins based on HMMSEARCH - belong to Chlamydia species, including almost 200 hits against non-bacterial proteins. In contrast, the HMM we derived here is almost exclusive to Chlamydia, matches a larger number of Chlamydia proteins (675 out of 744 Uniprot proteins), and identifies more than double the number of repeats per protein (average 6.1 repeats per protein, versus 2.9 for PF02415).

### Structural modeling shows Pmps share many features with other autotransporters

We took advantage of the newly released RoseTTAFold algorithm (38) to generate protein structure predictions for all *C. trachomatis* Pmps. RoseTTAFold claims accuracy approaching that of DeepMind’s record breaking Alphafold2 (40), is currently the top ranked method benchmarked by CAMEO (54), and has been made available by the Baker lab on their public Robetta protein structure prediction service (https://robetta.bakerlab.org/). In contrast, structure prediction with the older TrRosetta algorithm (38) results in less well resolved passenger domains, while the structures predicted by AlphaFold2 (40) for the homologous Serovar D proteins often show large completely unstructured loops (Figure 3). Because of the better detail in the side loops we decided to focus primarily on the RoseTTAFold structure predictions.

**Figure 3:**
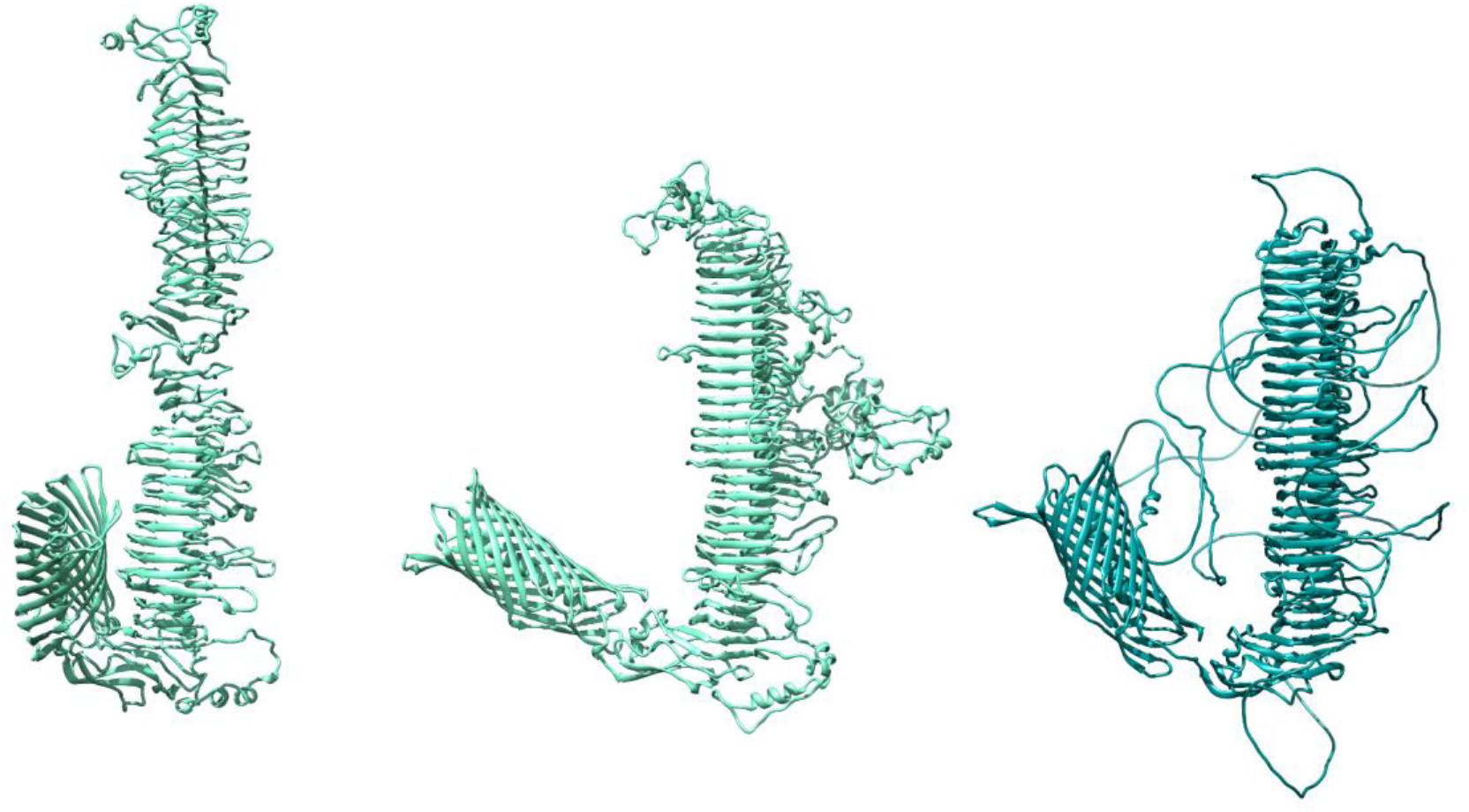
Comparison of structure predictions using TrRosetta, RoseTTAFold, and AlphaFold2. (A) TrRosetta structure of PmpC Serovar E. (B) RosetTTAFold structure of PmpC Serovar E. (C) AlphaFold structure of PmpC Serovar D.

The resulting structures show many of the stereotypical features found in other autotransporters, including a 12-stranded β-barrel at the C-terminal. The predicted β-barrel tended to incorporate multiple predicted mortise-tenon joints (55), and a β-hairpin structure as part of the fifth extracellular loop of the β-barrel that has been shown to be important for correct folding of the passenger domain in other autotransporters (56) (Figure 4).

**Figure 4:**
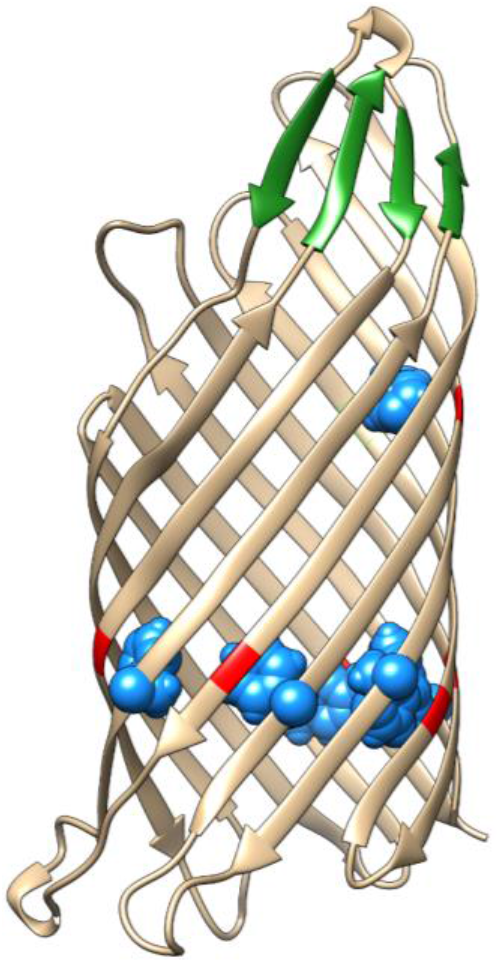
The β-barrel for PmpG showing six”mortise-tenon joints”, in which an aromatic side chain facing into the lumen of the barrel domain locks into a void created by the presence of a glycine residue on the neighboring β-strand. Blue: aromatic side chains. Red: glycine residues. Also shown is a 4-stranded β-sheet formed by β-hairpins in extracellular loops L4 and L5. The β-hairpin in L5 has been shown to be important for correct folding of the passenger domain in other autotransporters.

The conserved Pmp_M “Middle domain” is predicted to include the first couple of coils of the β-helix (even though it does not contain the conserved GGA(I,L,V)/FxxN motifs or HMM sequence pattern described above) that are most proximal to the β-barrel. The Pmp_M “Middle domain” flattens to a β-sandwich and is capped by a β-hairpin. This structure is similar to other autochaperone domains found in many autotransporters with β-helical passenger domains such as pertactin (57–59), suggesting it may play a role in proper secretion and folding.

As has been noted by others, the passenger domain is predicted to fold into a tightly coiled β-helix formed by parallel β-strands (Figure 5A). Based on the structure predictions shown here, Pmps and their homologs in related Chlamydia species would be some of the longest known parallel β-helix proteins, with at least 27 complete coils for PmpB and 22 coils for PmpD versus 16 coils for Pertactin.

**Figure 5:**
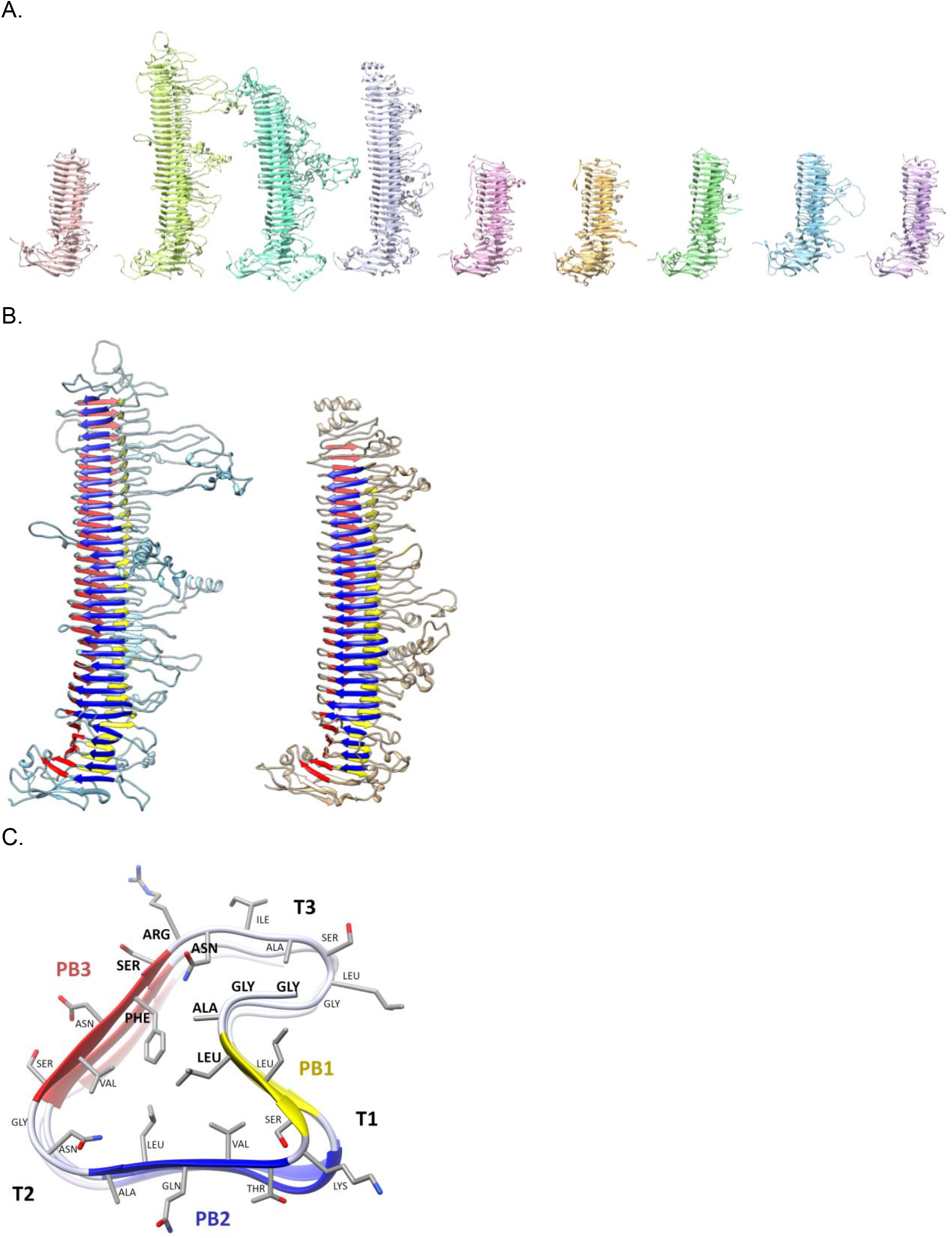
Pmp passenger domain structural models and nomenclature. (A) Family portrait of Pmp passenger domain models. From left to right: PmpA, B, C, D, E, F, G, H, and I. (B) Left: PmpD passenger domain showing 22 complete β-helix coils. Right: PmpD passenger domain cross section between amino acid residues 615-641 shows how the repeat sequence forms the core of the β-helix. The nomenclature follows Yoder et al., 1993 and Jenkins et al., 2001 (59,60). The parallel β-sheet PB1 is shown in yellow, PB2 in blue, PB3 in red. The GGA(I,L,V) and FxxN motifs are bolded-in this case we have GLY-GLY-ALA-LEU at the transition from T3 to PB1, and PHE-SER-ARG-ASN at the transition from PB3 to T3.

Following the nomenclature introduced by Yoder et al. 1993 and Jenkins et al. 2001 (59,60), the three parallel β-sheets (PB) in PmpD are labeled PB1 (yellow), PB2 (blue) and PB3 (red) and the turns (T) following the β-sheets labeled T1, T2, T3 (Figure 5C). Compared with the HMM pattern in Figure 2C, the initial conserved GGA constitutes the end of T3, followed by a short ~3 amino acid PB1 (Figure 3B). The two areas with sequence length variability between positions 6-9 and 16-17 (indicated by the vertical lines in Figure 2C) correspond to turns T1 and T2, while the conserved FxxN motif marks the transition from PB3 to T3 (Figure 3B). Note that the side chain positions of the most conserved amino acids in the GGA(I,L,V) and FxxN tetrapeptide motifs point inward, within the β-helix core (Figure 5C). These results suggest the tetrapeptide motif provide a more structural role to the passenger domain β-helix. Most of the variability in this pattern lies with the sometimes very long side loops inserted in T3 and occasionally T1, while T2 typically only varies in length by one or two amino acids (see also Figure 2C).

All predicted Pmp structures showed a 12-stranded β-barrel at the C-terminal, connected to the passenger domain by an α-helix threaded through the center of the β-barrel (Figure 4). Surprisingly, most models favored an angle of 90 degrees or more between the β-helical passenger domain and the transmembrane β-barrel, rather than having the two domains in a straight line. Coarse-grained molecular dynamics simulations suggest a flexible hinge region between the membrane-embedded β-barrel and the base of the passenger domain, allowing the extracellular portion of the protein to take on a range of conformations, from standing straight out from the membrane to lying flat against it, as illustrated in Figure 6.

**Figure 6:**
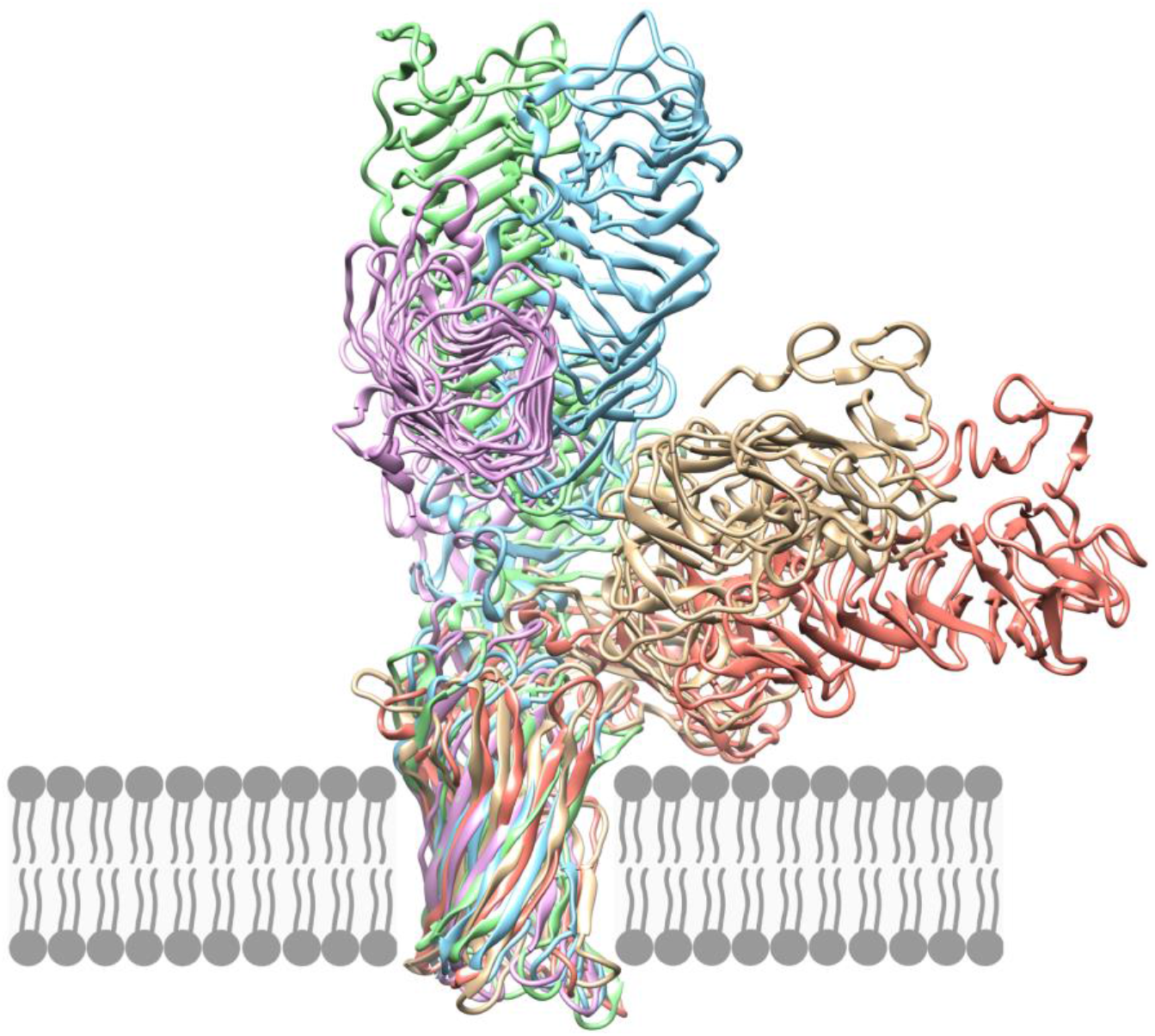
Coarse-grained molecular dynamics simulation of PmpE with the β-barrel domain embedded in a POPC bilayer shows that the passenger domain is connected by a flexible hinge between the β-barrel and the Middle domain.

### Protease cleavage sites are concentrated in side loops

Favaroni 2017 lists a total of 21 Pmp cleavage sites that have been experimentally detected in serovar L2 Pmp proteins by N-terminal sequencing (33), mass spectrometry (36), and identification of semitryptic peptides (25). Mapping these cleavage sites to our Serovar E structures, we found that three of these sites map to signal peptide cleavage sites, two cleave the alpha helix inside the transmembrane β-barrel (PmpD and PmpG), and two cleave the β-barrel itself. Of the remaining 14 cleavage sites that fall within the passenger domains, only two are located inside the β-helix coils, while the remaining 12 map to the side loops, the N-terminal cap, or the non-β-helical parts of the Middle domain.

### B-cell epitopes are concentrated in side loops, T-cell epitopes in the main β-helix of the passenger domain

The Immune Epitope Database (IEDB) (50) contains data on experimentally validated B-cell epitopes for 40 different *C. trachomatis* antigens, including 7 epitopes for PmpD and 5 for PmpC. All experimentally validated B-cell epitopes for PmpC and PmpD map to side loops off the β-helix or the Middle domain (Figure 9A). We also used Discotope 2.0 (52) to calculate B-cell epitope propensity scores based on the predicted structures for each Pmp, which also showed high-scoring regions predominantly in the passenger domain side-loops, and especially in some of the long extended loops (Figure 9B). The extracellular face of the transmembrane β-barrel also scores very high, even though IEDB shows no known epitopes in this area. The small extracellular β-barrel loops may be shielded from antibodies by the passenger domain, the chlamydial LPS, or neighboring membrane proteins.

**Figure 7:**
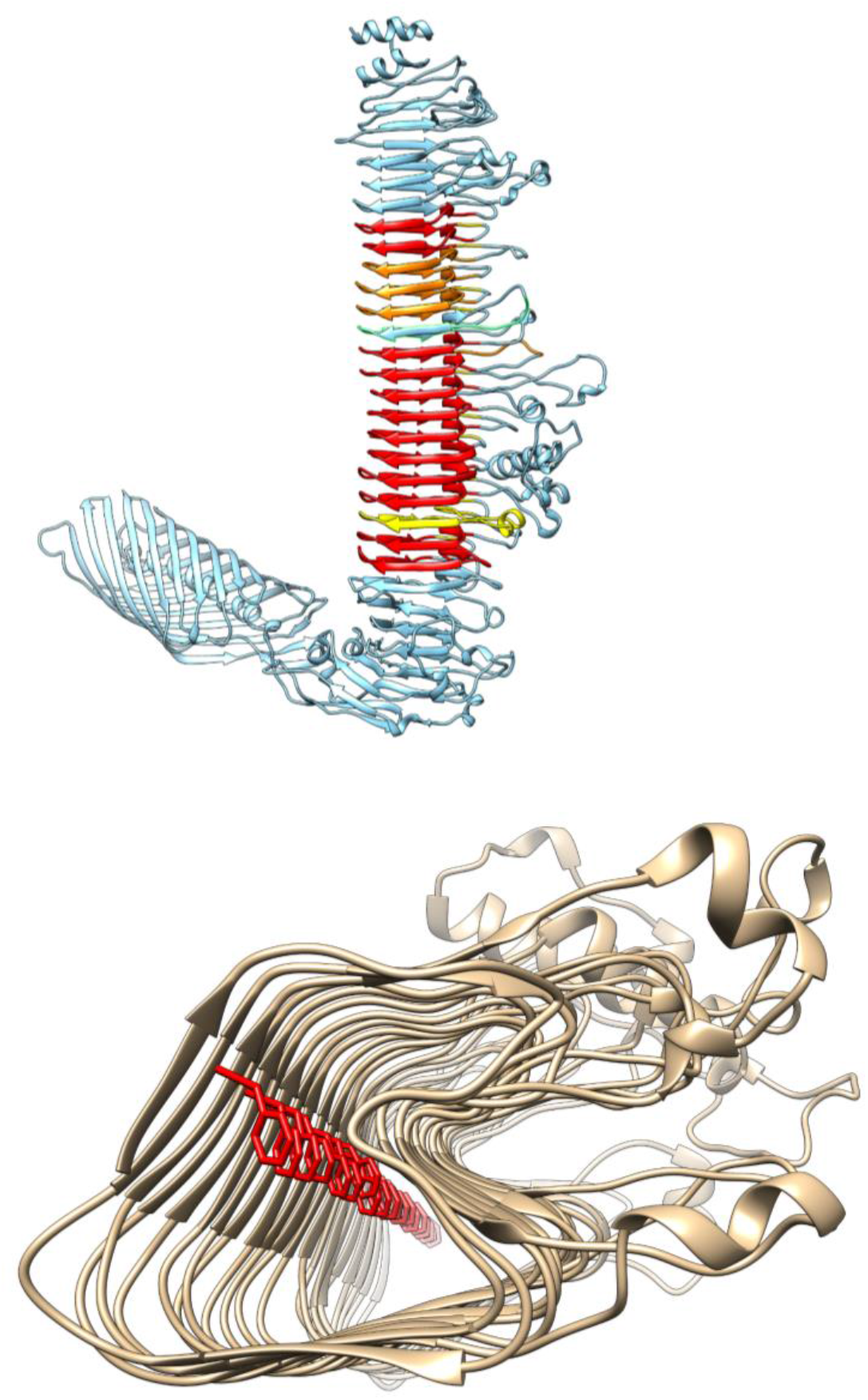
(A) Repeats on the PmpD structure. Repeats matching the core G-G-A-[ILV]-x(14,18)-F-x-x-N pattern are shown in red; those with a relaxed distance constraint or one aa mismatch from the core pattern are shown in orange; those matching only the more lenient HMM are shown in yellow. (B) alignment of PHE inside β-helix of PmpD.

**Figure 8:**
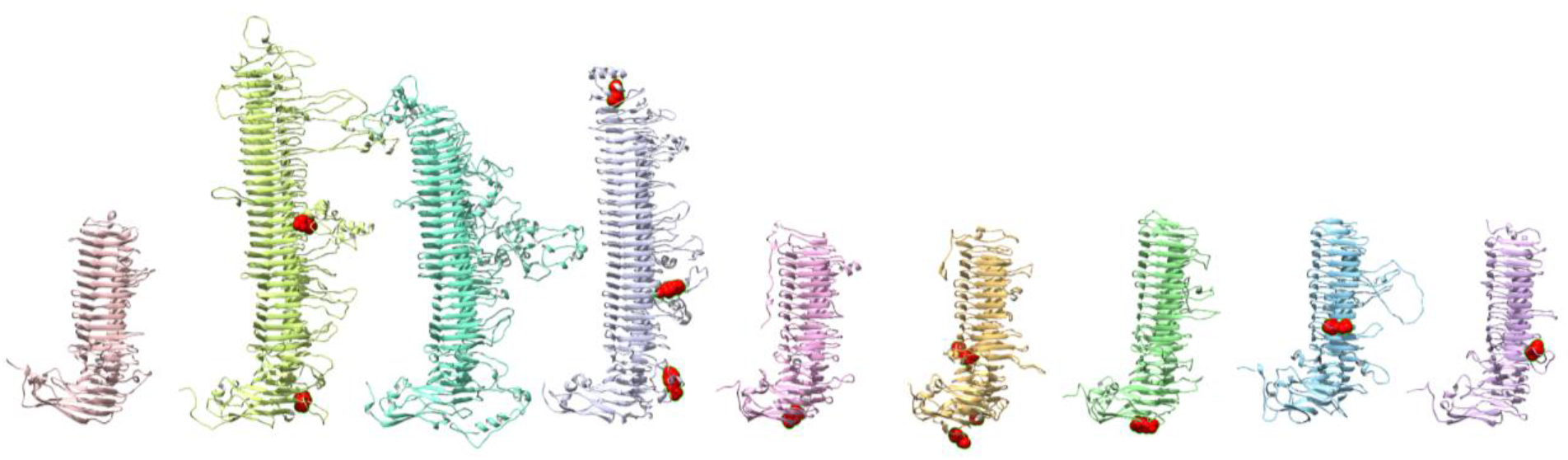
Pmp cleavage sites for serovar L2 Pmp proteins mapped to the serovar E passenger domain structures

**Figure 9:**
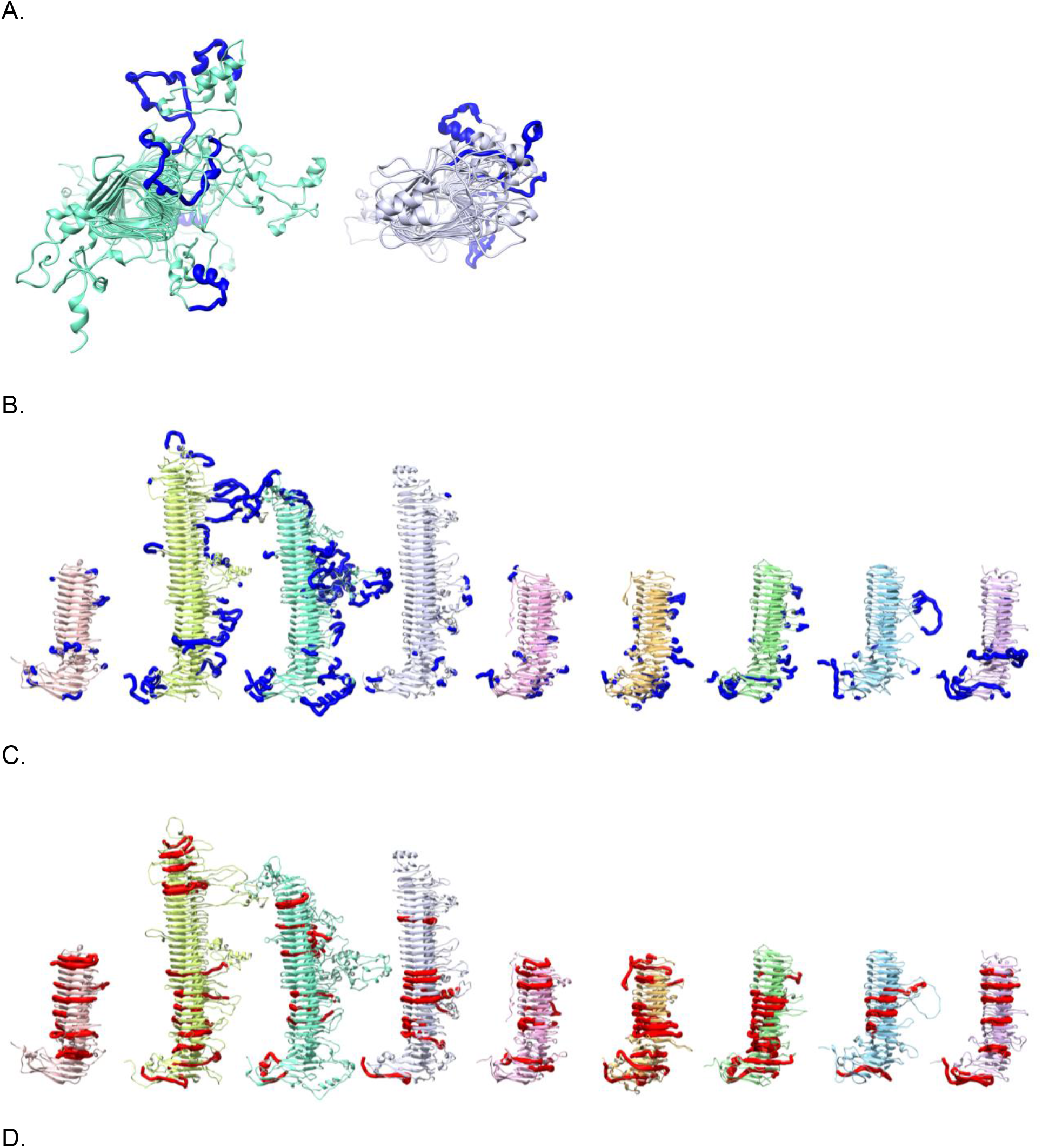
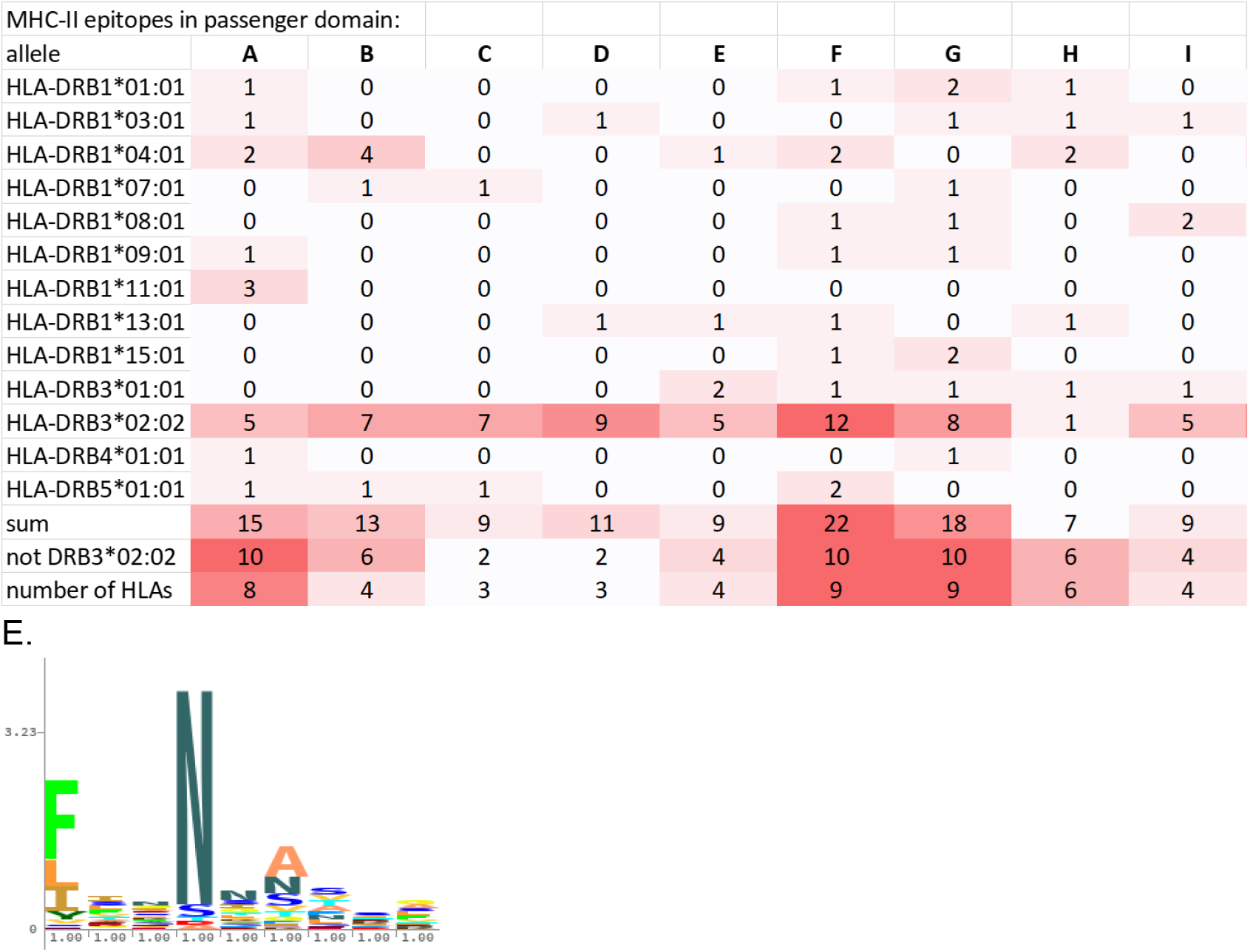
B-cell and T-cell epitopes on Pmp structure (A)Top view of the PmpC and PmpD passenger domains, with experimentally validated B-cell (B)Computationally predicted B-cell epitopes based on structure, using Discotope. From left to right: Pmp A, B, C, D, E, F, G, H, and I. (C)Computationally predicted MHC-II T-cell epitopes based on structure (D)The number of MHC-II T-cell epitopes differs considerably across Pmps and HLA subtypes (E)The core binding motif for MHC-II allele HLA-DRB3*02:02 overlaps with the FxxN repeat of the β-helix

Experimentally validated T-cell epitopes in IEDB occur for 49 different *C. trachomatis* antigens, which include one epitope each in PmpC, D, E, G, H, and I. Two MHC-I epitopes map to different strands of β-barrel for PmpC and PmpI, while the four MHC-II epitopes in PmpD, E, G, and H map to various locations along the β-helix.

Computational prediction of MHC-II T-cell epitopes map predominantly to the β-helix of the passenger domains (Figure 9C), with some epitopes matching multiple HLA subtypes. Interestingly, epitopes for allele HLA-DRB3*02:02 seem strongly overrepresented, representing more than one third of all MHC-II T-cell epitopes (Figure 9D). Upon further examination, it appears that the core binding motif for this allele overlaps with the FxxN motif in the passenger domain β-helices (Figure 9E). This suggests that populations with this particular MHC-II allele may be able to mount a stronger T-cell immune response to Chlamydial infections. The number of MHC-II T-cell epitopes differs noticeable across Pmps, with the small PmpA containing 25 epitopes (20 not counting HLA-DRB3*02:02) for 12 out of the 13 HLA subtypes tested, while the equally short PmpH only contains 9 (8 not counting HLA-DRB3*02:02), and the much longer PmpC contains 13 (6 not counting HLA-DRB3*02:02) (Figure 9D). This suggests that some Pmps may present better vaccine targets to stimulate a broad T-cell response. MHC-I T-cell epitopes were predicted to be fairly abundant in Pmps, and spread throughout the length of the entire protein.

### Side loops are predicted to be involved in host cell adhesion

Pmp side loops include proline-rich regions (PRRs) that may be involved in binding to host membrane proteins. PRRs are often involved in eukaryotic protein-protein interactions, and can be exploited by viral and bacterial pathogens to interact with host proteins (61,62). In the Pmps, PRRs are predominantly found on the side loops and the Middle domain (Figure 10)

**Figure 10:**
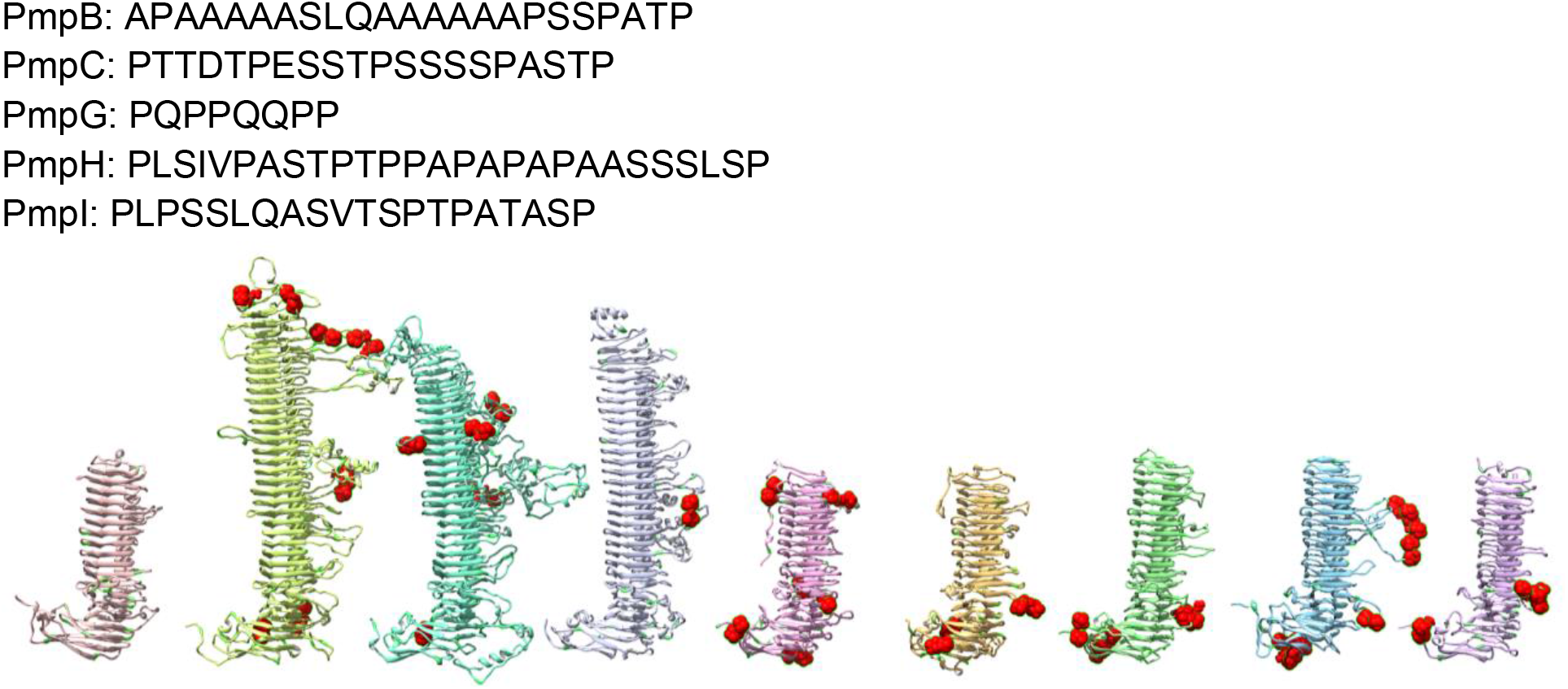
Proline-rich regions are located primarily in the side loops and the Middle domain. Sites with at least two prolines separated by 0-2 residues are highlighted in red.

Besides binding to membrane proteins, Pmps may also interact directly with the host membrane. As expected, prediction of protein-membrane interface peptides using DREAMM yields multiple hits in the transmembrane beta barrel for each Pmp. However each Pmp also has one or more predicted membrane-penetrating amino acids at the N terminal and in various side loops off the main beta helix, that could be involved in adhesion to host membrane (Figure 11).

**Figure 11:**
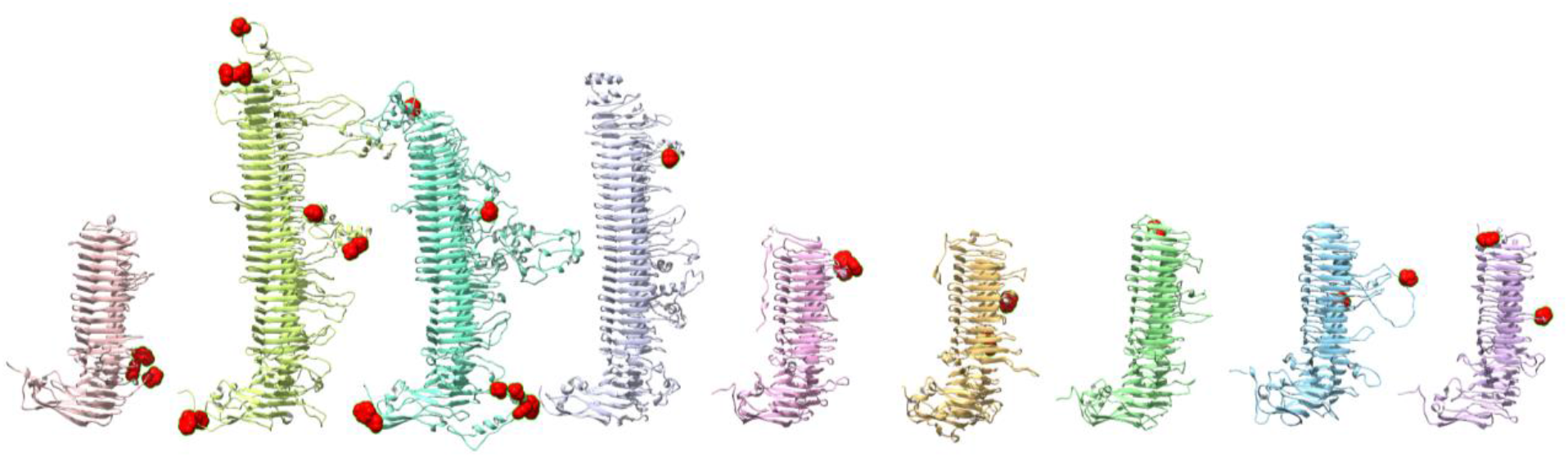
Predicted membrane-penetrating amino acids (highlighted in red) are located primarily in the side loops, and at the N terminal, as well as the transmembrane beta barrel (not shown).

Sequence variation between the 15 *C. trachomatis* serovars, based on a multiple sequence alignment with MAFFT is also more concentrated in the passenger domain side loops (Figure 12), which may be correlated both with positive selective pressure on the B-cell epitopes by the host immune system, and with differences in tissue tropism.

**Figure 12:**
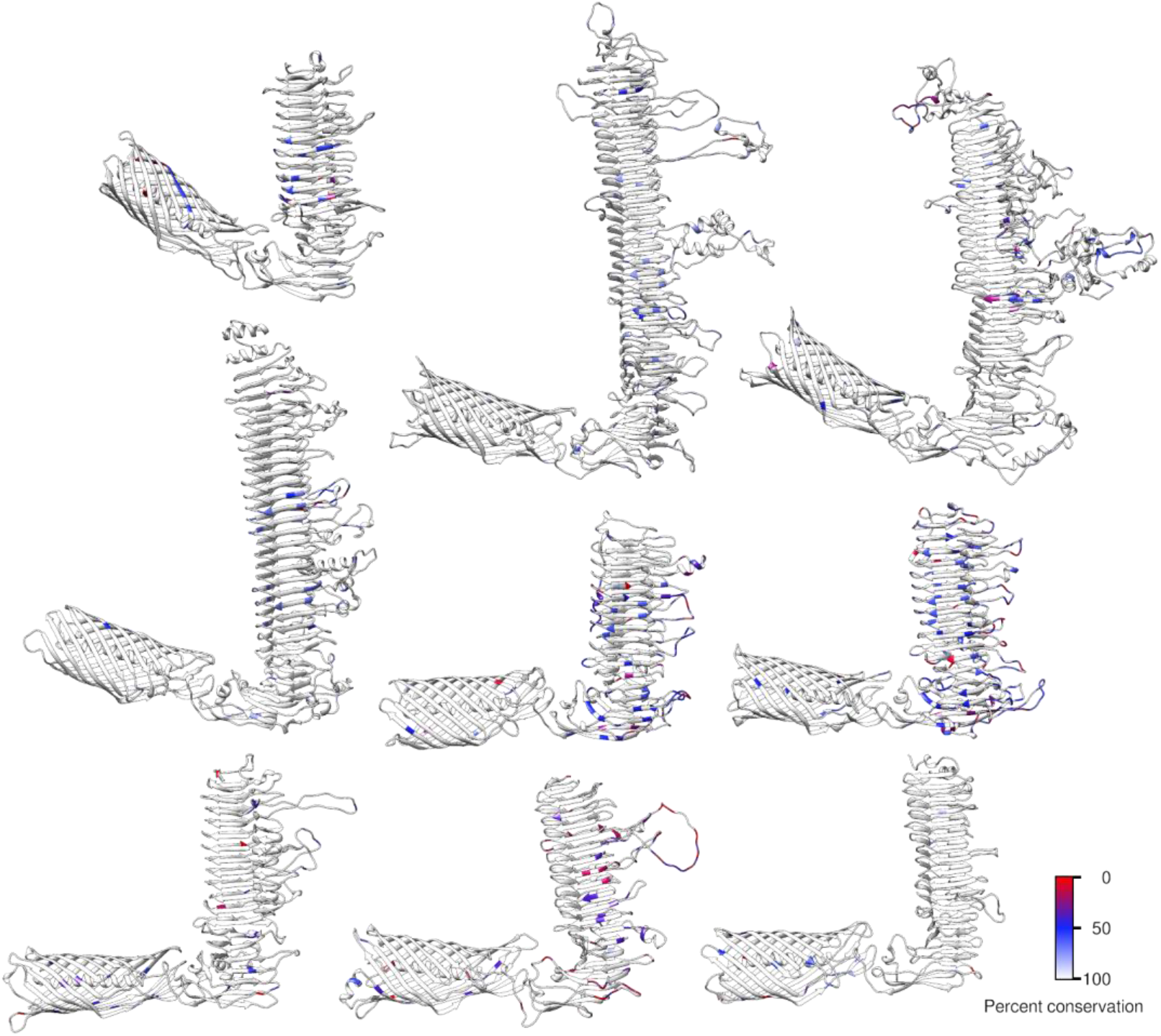
Conserved Pmp amino acids sequences within *C. trachomatis*. From left to right: Top row) Pmp A, B, and C. Middle row) Pmp D, E, and F. Bottom row) Pmp G, H, and I. Amino acid sequences from one Pmp were compared to the same Pmp across the main 15 *C. trachomatis* serovars by MAFFT.

## Discussion

The Pmp family is a group of surface exposed proteins in *Chlamydia* that are viable vaccine candidates. These proteins are naturally immunogenic in humans and N-terminal subunit vaccines offer protection to *Chlamydia* challenges in non-human primates and mice. However, in lieu of a crystal structure, few resources have utilized predictive protein structural modeling to characterize and map features within the Pmp family. In this study, we took advantage of the newly released RoseTTAFold algorithm to create full-length protein structure predictions for all 9 Pmps in *C. trachomatis* Serovar E (Bour). Our bioinformatic and structural analyses discovered the GGA(I,L,V) and FxxN tetrapeptide motifs are part of a larger repeat structure within the passenger domain β-helix core. Moreover, we found the passenger domain side loops that jut out of the β-helix contain many interesting features - including protease cleavage, host cell adhesion, and B-cell epitopes; while T-cell epitopes are predominantly found in the β-helix itself. These results highlight the modular structure of Pmp passenger domains that may lend itself well to protein engineering and rational vaccine design.

Although the GGA(I,L,V) and FxxN tetrapeptide motifs have been recognized as prominent features of the Pmp passenger domains for more than twenty years (16), their semi-regular spacing had not been previously appreciated. Here we have shown that the tetrapeptide motifs fit into a larger repeat sequence that can be fit with an HMM model including several other partially conserved residues, and that these larger sequence repeats correspond to the individual coils of the predicted β-helical structure of the passenger domains. Since not all the β-helical coils include the canonical GGA(I,L,V) and FxxN motifs, it may be possible to derive a more inclusive HMM by starting from a structural alignment of the coils instead.

Cartoon representations of autotransporters typically show the passenger domain in line with the transmembrane beta barrel sticking out from the membrane at a right angle (58,63–66).However structural modeling of the Pmps shows a variety of angles between the beta barrel and the passenger domain, and molecular dynamics shows the presence of a flexible hinge between the two domains. It is not clear whether this is unique to the Pmp family, or a more general feature of autotransporters. It should be noted that as of yet no full-length crystal structures exist of Pmp proteins, or indeed any autotransportes with β-helical passenger domain. The potential for passenger domains to lay flat against the outer membrane also opens the possibility for Pmp’s to make longer-range protein-protein interactions with other members of the Outer Membrane Complex.

Mutating or deleting the conserved GGA(I,L,V) and FxxN motifs has been shown to disrupt Pmp oligomerization (21) and host cell adhesion (21,67). Our structural models clearly show that the conserved residues of the tetrapeptide motifs are actually located on the inside of the beta helix (see Figure 5C). This suggests that rather than being directly involved in binding to host membranes, host proteins, or other Pmp proteins, these motifs are more likely to play a purely structural role, ensuring proper folding and stacking of the coils of the β-helical passenger domain.

This enhanced understanding of the structural role of the tetranucleotide motifs shifts the perspective towards the areas between these conserved motifs. Indeed, we have shown that the side loops jutting out from the β-helical backbone of the passenger domain seem to play an important role in a variety of processes, including protease cleavage, antibody binding, and host cell adhesion.

A recent Genome-Wide Association study of *C. trachomatis* serovar G genomes identified polymorphisms in Pmp E, F, and H that are associated with rectal tissue tropism, localized specifically to the side loops in these Pmps (68). Pmps are also known to be directly involved in invasion of host cells by binding to host EGFR in *C. pneumoniae* (69) and C. psittaci (70). Although the precise mechanisms of these interactions are as yet poorly understood, the proline-rich regions and membrane interaction domains we identified in the side loops may be involved in mediating these interactions. PmpD also contains an integrin-binding RGD motif on a short loop in turn T1, that may be involved in attachment or entry (33)

The ideal *Chlamydia* vaccine would provide cross-serovar protection with strong cell-mediated and humoral responses. Russi et al. (71) had previously analyzed T-and B-cell epitopes in PmpD. Here we have extended that analysis to the entire Pmp family. We are also leveraging much higher quality structure predictions than they had available at the time, which should especially benefit the B-cell epitope prediction. For example, several of the sequence-based B-cell epitope predictions they derived using Bepipred (51) overlapped with the tetranucleotide motifs, which we predict are embedded in the beta helix and therefore unlikely to be accessible to antibodies. They decided to focus on six peptides that included multiple B-and T-cell epitope predictions. However our analysis shows that B-and T-cell epitopes are largely disjoint, with B-cell epitopes predominantly in side loops, while MHC-II T-cell epitopes are concentrated more inside the beta helix itself. For vaccine design it may therefore be more effective to choose B-and T-cell epitopes separately, rather than expect to be able to capture both with the same peptides.

## Acknowledgements and Funding

This work was performed, in part, under the auspices of the U.S. Department of Energy by Lawrence Livermore National Laboratory under Contract DE-AC52-07NA27344. Support was also provided by NIH grant U19AI144184.

